# An approach to rapid distributed manufacturing of broad spectrum anti-viral griffithsin using cell-free systems to mitigate pandemics

**DOI:** 10.1101/2022.12.19.521044

**Authors:** Shayan G. Borhani, Max Z. Levine, Lauren H. Krumpe, Jennifer Wilson, Curtis J. Henrich, Barry R. O’Keefe, Donald Lo, G. Sitta Sittampalam, Alexander G. Godfrey, R. Dwayne Lunsford, Venkata Mangalampalli, Dingyin Tao, Christopher A. LeClair, Aaron Thole, Douglas Frey, James Swartz, Govind Rao

## Abstract

This study describes the cell-free biomanufacturing of a broad-spectrum antiviral protein, griffithsin (GRFT) such that it can be produced with consistent purity and potency in less than 24 hours. We demonstrate GRFT production using two independent cell-free systems, one plant and one microbial. Griffithsin purity and quality were verified using standard regulatory metrics. Efficacy was demonstrated *in vitro* against SARS-CoV-2 and HIV-1 and was nearly identical to that of GRFT expressed *in vivo*. The proposed production process is efficient and can be readily scaled up and deployed anywhere in the world where a viral pathogen might emerge. The current emergence of viral variants has resulted in frequent updating of existing vaccines and loss of efficacy for front-line monoclonal antibody therapies. Proteins such as GRFT with its efficacious and broad virus neutralizing capability provide a compelling pandemic mitigation strategy to promptly suppress viral emergence at the source of an outbreak.

## Introduction

Currently, manufacturing a drug or vaccine is done in large and expensive facilities that have been inspected and approved by regulatory authorities^1^. The drug substance typically requires extensive testing and characterization for purity and potency and must go through extensive clinical safety and efficacy trials. Manufacturing facilities are centrally located and rely on well-developed transportation networks for distribution of the product usually via a cold chain. During the early phases of pandemic progression, before vaccines and mAbs can be developed and administered, there is a gap in our current prevention strategies. As recent experience indicates, that gap cannot be effectively filled by masking and physical distancing (both of which encounter significant societal resistance)^2–4^.

The COVID-19 pandemic has highlighted transformational advantages offered by broad-spectrum antiviral drugs that both prevent and treat viral infections at the early stages of an outbreak ^5^. Initial enthusiasm for such compounds such as hydroxychloroquine^6^, ivermectin^7^, and oleandrin^8^ was shown to be misplaced^9–11^, and the need for effective agents to fill a public-health gap early in an emerging pandemic remains. Additionally, promising therapies such as remdesivir, baricitinib, and dexamethasone, which were widely used in treating critically-ill COVID-19 patients, proved efficacious only in a limited number of cases^12^. Modern travel and international trade accelerate viral transmission making effective control nearly impossible with current measures. This suggests that a strategy that can mitigate viral transmission early in the initial outbreak, before more targeted vaccines and antivirals can be available, would be extremely valuable in reducing the human cost of future pandemics. While biologics such as monoclonal antibodies (mAbs) have been shown to be viable methods for the treatment of COVID-19, these drugs are slow and costly to produce and are specifically targeted^13–15^. The rapid onset of resistance to various mAb and antiviral treatments in emerging variants of SARS-CoV-2^16,17^ further highlights the need for easily produced antiviral agents with a mechanisms of action equally effective on a broad range of emerging viral variants.

In this paper, we report the development of such a system for the rapid delivery of the broadspectrum antiviral protein, Griffithsin (GRFT), using both bacterial and plant-based cell-free protein synthesis (CFPS) complemented with efficient purification technology. GRFT, a 12.7 kDa protein originally isolated from a red alga has affinity for high-mannose oligosaccharide moieties on the surface glycoproteins of enveloped viruses and has been shown to inhibit viral entry into host cells^18^. This mechanism of action assures efficacy even against mutant viruses, unlike current vaccines and antivirals. GRFT is an ideal candidate for this project as exerts activity against all known *Coronaviridae*, including SARS-CoV-1^19^, MERS^20^, and SARS-CoV-2^21,22^. GRFT also provides demonstrated *in vivo* efficacy against other viral families including *Paramyxoviridae*^23^, *Flaviviridae*^24^, *Retroviridae*^25^, and *Hantaviridae*^26^. Further, GRFT is: readily produced in highly productive recombinant systems^19,27,28^, stable at high temperatures and low pH^18^, and resistant to protease digestion^29^. It has also been shown to be non-toxic and non-immunogenic in multiple animal model systems^30–32^. Finally, GRFT and its oxidation-resistant analog Q-GRFT^33^ have both progressed to Phase I clinical trials in humans using multiple routes of administration, including intranasally, for prevention of SARS-CoV-2 infection with little evidence of immunogenicity^34,35^.

The COVID-19 pandemic is but one serious example of the zoonotic transmission of viral disease. We have now all experienced the human and economic costs of limited public health options during an outbreak of such an aerosol-transmissible virus. To limit the societal risks arising from future zoonotic pathogens, we propose the development of point-of-care (POC), on-demand production systems for a previously validated broad-spectrum antiviral agent. Portable, on-demand drug manufacturing platforms^36–38^ can produce antiviral proteins on the time scale of hours, making them ideal for combating future pandemics. To build a more durable and resilient antiviral supply chain, we propose a portable, highly productive technology based upon the proof of concept described here. A rapidly implemented and portable cell-free biomanufacturing platform (Figure 1) would provide GRFT to suppress potential outbreaks close to their points of origin^39,40^. Importantly, we demonstrate an approach that goes beyond first observational studies and includes methods and metrics for achieving the final product purity, identity and potency that will be required by regulatory authorities. We also provide a process technology foundation for efficient, POC manufacturing.

**Figure 1.**
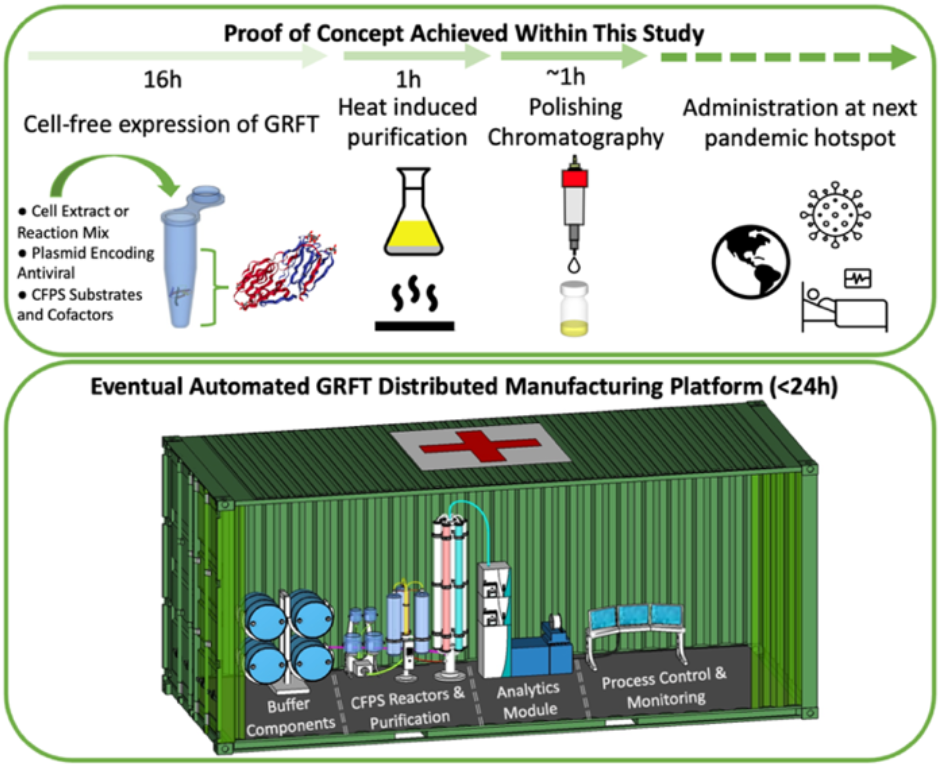
Top: Proof of concept for GRFT bioprocess (top) includes expression using a cell-free system (CFS), heat induced purification and polishing chromatography to yield active and pure GRFT for use at the next pandemic hotspot. Bottom: A distributed manufacturing platform (bottom) which can be deployed in a shipping container to make biologics safely and effectively at the site of the future viral outbreak. The container includes concentrated, dried solutions for reconstitution of cell-extracts, required CFPS reagents, and purification buffers, as well as validated analytics for real-time process monitoring and control.

## Materials and Methods

### Griffithsin Gene Construction and DNA preparation

A tagged wild-type GRFT gene, tagged-GRFT (Supplementary Figure 1a), was designed (Genscript) for production of an N-terminal hexahistidine tagged GRFT with an enterokinase cleavage site after the tag (14.5 kDa). The gene was subcloned via T4 DNA ligation into the ALiCE-01 expression vector, for expression in ALiCE. Additionally, two versions of the tagless M78Q-GRFT gene, tagless-GRFT.std (12.68 kDa), were introduced through Gibson Assembly between a T7 promoter and T7 terminator in a pUC19-based vector (NEB), pMZL, with varying codon optimization strategies. The first version, tagless-GRFT.std, was comprised of the most frequently used *E. coli* codons (Genscript). A second version, tagless-GRFT.opt (12.68 kDa) (Supplementary Figure 1a), was synthesized by IDTdna with the most common *E. coli* codons except for those encoding glycine and serine. These codons comprise 18.2% and 15.7% of the amino acid composition respectively, of the coding sequence and were randomly chosen to minimize codon replication. In addition, the glycine and serine codon choice was then adjusted to minimize mRNA secondary structure while preserving the encoded amino acid sequence. The AGC serine codon was avoided. The resulting amino acid composition for both constructs is provided in Supplementary Figure 1b. DNA plasmid preparation for CFPS reactions was conducted using the Qiagen Maxiprep kit (Cat# 12162) using the recommended protocol with water as the elution buffer.

### *E. coli* Cell Extract Preparation

KC6 *E. coli* cells^41^ were produced by continuous culture at a growth rate of 0.8 hr^-1^ and an OD_660_ of about 35 using an Eppendorf Bioflo320 10 L bioreactor. A total of 40 L of defined media was used. The cells exiting the bioreactor were immediately cooled as the exit stream passed through a copper cooling coil immersed in an ice bath. The broth was centrifuged at 5000 x g for 15 minutes, and the cell paste was frozen into sheets. The next day, the sheets were broken into pieces and resuspended in 1 ml of S30 buffer (10 mM Tris-Acetate pH 8.2, 14 mM Mg Acetate, 60 mM K Acetate) per gram of cell paste within a 20L and mixed using a motorized impeller until no visible chunks were seen. The resuspension was thawed quickly using 30°C water circulated through an immersed coil while keeping the cell suspension temperature below 8°C. The resuspended cell mass was lysed by a single pass through a high-pressure homogenizer (EmulsiFlex-C5, Avestin, Canada) at 17,500–25,000 psi. The resulting homogenate was clarified by a 5000 x g, 30 min centrifugation at 4 °C. The resulting pellets were discarded, and the clarified cell extract was frozen. The extract was then thawed and incubated at 37°C for 60 min for full activation. A final 20 min centrifugation step at 5000 x g was performed and the resulting cell extracts were flash-frozen and stored at −80 °C.

### *E. coli* Cell-free Expression and Quantification

*E. coli* CFPS of GRFT was conducted at a 25 μL scale in 2 ml Eppendorf tubes using the PANOx-SP system utilizing a phosphoenolpyruvate (Roche, Indianapolis, IN), amino acids (Sigma Aldrich, St. Louis, MI), nicotinamide adenine dinucleotide (Sigma Aldrich), oxalic acid (Sigma Aldrich), spermidine (Sigma Aldrich), and putrescine (Sigma Aldrich) mastermix solution as described previously^42^ with the following modifications: 15.6 ng/μL pMZL-tagless-GRFT plasmid, addition of 50 mM HEPES, 2 mM of all amino acids with tyrosine prepared separately due to insolubility within the mixture, and the addition of 5 μM L-[^14^C(U)]-leucine (PerkinElmer, Waltham, MA). All reactions were performed in triplicate at 30 °C for approximately 16hrs. Quantification of synthesized GRFT was determined based on incorporation of ^14^C-leucine into TCA-precipitable protein using a liquid scintillation counter to measure incorporated radioactivity. Soluble versus insoluble protein yields were determined as previously reported^43^.

### Autoradiogram Preparation

*E. coli* CFPS samples containing radiolabeled GRFT were diluted in NuPAGE™ LDS Sample Buffer with 80 mM DTT. The solution was subsequently incubated at 95 °C for 10 min to completely denature GRFT. The samples were loaded onto a 4-12% SDS-PAGE gel (Invitrogen, Waltham, MA) and run using the MES SDS running buffer (Invitrogen). The gels were dried and contacted with a storage phosphor screen overnight (Molecular Dynamics, Sunnyvale, CA). Autoradiograms were visualized using a Typhoon imaging system (GE Healthcare, Uppsala, Sweden).

### ALiCE^®^ Expression

A commercially available plant-based CFPS kit (ALiCE^®^), utilizing tobacco BY-2 (*N. tabacum*) cell extracts, was donated by LenioBio (Aachen, Germany). The proprietary cell-free reaction mixture contains all factors required for *in vitro* transcription (e.g., RNA polymerase and NTPs) and translation (ribosomes, translation/initiation factors, tRNAs, etc.). Two DNA vectors were available for subcloning of our tagged-GRFT construct, pALiCE01 and pALiCE02, for cytosolic and microsomal expression, respectively. Given the absence of extensive post-translational modifications in GRFT we opted to express the tagged-GRFT construct in the pALiCE01 vector for protein expression in the cytosol. The CFPS reaction was conducted at a 50 μL to 500 μL scale, by mixing lysate and plasmid DNA (pALiCE01 tagged-GRFT) and distribution into 96, 48, and 24-well flat bottom plates (Corning, Cambridge, MA), respectively. Reactions were performed in triplicates and completed reaction solutions were pooled for further analysis. The wells surrounding the reaction wells were filled with nuclease-free water to prevent loss of reaction volume. The reaction was then incubated at 25 °C and 500 rpm for 16-18 h in an Eppendorf Thermomixer C. For determination of soluble protein concentration, the supernatant of post expression samples was collected after centrifugation at 10,000 x g for 10 min. Quantification of synthesized GRFT was based on Western blot analysis by comparison to gel bands with known amounts of product.

### Western Blot Analysis

In a fresh 1.5 mL Eppendorf tube, 1-29 μL of 1X PBS was aliquoted and mixed with 1-29 μL of the sample. Samples were then treated with 2-6 μL of 6X Tris-glycine sample buffer, then boiled at 100 °C for 5 min. Samples were spun and then loaded onto a 4–20% Criterion™ TGX gel (Bio-Rad, Hercules, CA) and run at 250 V for 30 min. The gel cassette was then opened, and the gel was transferred into the blotting apparatus immersed in 1X Tris-glycine Transfer buffer. A nitrocellulose membrane which was submerged in Transfer buffer for 5 min was placed on the gel apparatus, allowing for protein transfer onto the nitrocellulose membrane. Upon completion of protein transfer at 100 V for 1 hour, the membrane was submerged in 20 mL of blocking solution for 1 h. The blot was probed with mouse anti-GRFT antibody (provided by Barry O’Keefe, NCI) at a 1:1000 dilution for 1 h, followed by three washes (5 min each) of 1X PBS pH 7.4, 0.05% Tween-20 (PBS-T). The blot was then probed with goat anti-rabbit HRP-conjugated secondary antibody at a dilution of 1:5000 for 1 hour, washed, and then developed using SuperSignal West Dura (Thermo Fischer, Waltham, MA) and imaged using a Thermo Fisher Scientific myECL Imager. A standard curve ranging from 250 – 1000ng was generated using GRFT standard (1mg/mL) for qualitative and quantitative analysis of GRFT samples. The resulting bands in the standard curve lanes were quantified using ImageJ densitometry and used to calculate the concentration in the unknown samples.

### Coomassie and Silver Staining

Gel loading samples were prepared in a similar manner as mentioned in the Western blotting procedure. For silver staining, the samples were analyzed using a ProteoSilver plus silver stain kit (Sigma Aldrich, St. Louis, MI) and gel preparation were carried out according to kit protocol. For Coomassie stain, the gel was wash three times with deionized water, 5 min each, then immersed in Coomassie Brilliant Blue R-250 stain for 1h with gentle shaking. The gel was then de-stained using a buffer consisting of 50% deionized water, 40% methanol, and 10% acetic acid for a minimum of 2hrs, until the background became clear. The percent purity was calculated based on ImageJ densitometry, by taking the ratio of the lowest detectable GRFT band (peak area) versus the total peak area, where total peak area is equal to the lowest detectable GRFT band plus the peak area of impurities found in an overloaded gel lane. A standard curve ranging from 125 – 1000ng was generated using GRFT standard (1mg/mL) for qualitative and quantitative analysis of GRFT samples. The resulting bands from the standard curve were quantified using ImageJ densitometry and used to calculate the concentration of the unknown samples.

### Heat Precipitation of tagless and tagged GRFT

The unusual thermal and pH stability of GRFT enabled the protein to be a prime candidate for heat-induced precipitation^28,44,45^. Four factors have been previously reported to impact the yield and resultant purity of GRFT including: ammonium sulfate concentration in the GRFT solution, incubation temperature, pH, and incubation time. We selected the optimum conditions determined by the design of experiments previously reported by Decker et al^28^; a 1:1 dilution of the cell-free harvest in 20.5% (NH4)2SO4 (%sat), 60°C, pH 3.4, and 1 h, respectively. Adjustment of pH was achieved using concentrated HCl and NaOH. Assembly of precipitation mixture took place in 2 mL Eppendorf tubes with incubation in an Eppendorf Thermomixer C, without agitation. After 1 h incubation period, precipitated proteins were sedimented by 10,000 x g centrifugation for 5 min at room temperature. The supernatant fractions were then buffer exchanged into sterile 1X PBS (pH 7.4) for further analysis. Samples were then analyzed using sodium dodecyl sulfatepolyacrylamide gel electrophoresis (SDS-PAGE) Coomassie and Western blot analysis. Percent recovery was calculated by multiplying the concentration of each sample by the volume collected for each sample, then dividing by the total GRFT harvest mass.

### Immobilized Metal Affinity Chromatography (IMAC) capture of tagged-GRFT

Purification of tagged-GRFT was achieved by using a zinc-chelating resin (binding capacity 20mg/mL) (G-Biosciences). A calculated amount of resin was added to a 0.8 mL Pierce centrifuge spin column based on the estimated total protein yield obtained by Western blot analysis. Before protein binding, the resin was washed twice with 5 column volumes (CV) of distilled water and spun at 500 x g for 2 min. A binding buffer consisting of 50 mM phosphate buffer, 300 mM NaCl, 10 mM Imidazole, and pH 8.0 was used at a 1:1 ratio (binding buffer to harvest). The sample was then subject to mixing on a rotary shaker for 20 min at room temperature. Samples were then spun down and flow-through (FT) was collected for future calculation of recovery. Post-loading impurities were washed by adding 5 CV of binding buffer (W1) and spinning at 500 x g for 2 min. The sample was then eluted with 0.5 CV of an elution buffer containing 50 mM phosphate buffer, 300 mM NaCl, and 250 mM Imidazole, pH 8.0. Elution samples were then analyzed using sodium dodecyl sulfate-polyacrylamide gel electrophoresis (SDS-PAGE) Coomassie and Western blot analysis. Percent recovery was calculated by multiplying the concentration of each sample by the volume collected for each sample, then dividing by the total GRFT harvest mass.

### SEC Analysis

The homodimeric properties of GRFT were analyzed using an UltiMate™ 3000 HPLC instrument (Thermo Fischer Scientific, Rockford, IL) with a 125 Å 4.6 x 300 mm TSKgel SuperSW2000 column (Tosoh Biosciences, King of Prussia, PA) equilibrated with 1X PBS (pH 7.4) at a flow rate of 0.3 mL/min. 10-100 μL of the sample was injected into the column and aliquots (~100 μL) of the flow-through were collected. A protein mix standard (Sigma Aldrich, St. Louis, MI) containing thyroglobulin bovine (640 kDa), γ-globulins from bovine blood (150 kDa), albumin chicken egg grade VI (44 kDa), ribonuclease A type I-A from bovine pancreas (13.7 kDa) and p-aminobenzoic acid (pABA) (137 Da) was used. Absorbance was measured at 280 nm and the retention times (RT) obtained from the protein mix were then plotted on a semi-log plot and fitted with linear regression for the estimation of molecular weights (MW) for the various GRFT constructs.

### HPLC-TOF-MS analysis

Liquid chromatography (LC) was performed on an Agilent 1290 Infinity II LC system (Agilent Technologies, Wilmington, DE, U.S.A.) equipped with a diode array detector, binary pump, multicolumn thermostat, and autosampler. The mobile phases used for the separation were MS-grade water with 0.1% formic acid (solvent A) and MS-grade acetonitrile with 0.1% formic acid (solvent B). Gradient elution of proteins from the analytical column (HALO BioClass Protein Diphenyl, 1000 Å, 2.7 μm, 2.1 x 100 mm) was performed using a gradient starting at 5% B at a flow rate of 0.3 mL/min. The mobile phase was then increased linearly from 5 to 80% B over 8 min, and maintained at 80% B for 3 min. This was followed by 5% B for 5 min to re-equilibrate the column. Separations were performed at a column temperature of 35 °C. Sample injection volume was adjusted to improve analytical sensitivity. Mass spectrometry (MS) experiments were conducted on an Agilent 6230 TOF system (Agilent Technologies, Wilmington, DE, U.S.A.), equipped with a DUAL AJS ESI source operating in positive ion mode. MS spectra were acquired from m/z 400 to 3200 at a scan rate of 1 spectrum per second with Profile format. The applied electrospray ionization (ESI) source parameters were as follows: gas temperature, 300 °C; gas flow, 7 L/min; nebulizer, 35 psi; sheath gas temperature, 275 °C; Vcap, 4500 V; nozzle, 1000 V; fragmentor, 275 V. HPLC-TOF data analysis was performed with Agilent MassHunter Qualitative Analysis (B.07.00) utilizing the BioConfirm software (B.07.00) workflow for intact protein analysis. Protein deconvolution employed the following parameters: deconvolution algorithm, pMod; mass range, 4,000 to 50,000 (mass step at 1.0 Da); baseline factor, 7.0; adduct, Proton.

### Anion Exchange Chromatography and Endotoxin Removal

Anion exchange (AEX) chromatography was performed for the removal of bacterial endotoxin in a similar manner to Chen et al.^46^ using a 1 mL HiTrap™ Q FF column (Cytiva, Marlborough, MA) in flow-through mode. After equilibrating the column with 100 mM citrate buffer at pH 3.0, 125 μL of the sample was loaded onto the column. and chased with equilibration buffer at 1mL/min. where 250uL fractions were collected. Quantification of bacterial lipopolysaccharides (LPS) was achieved by using the Pierce Chromogenic Endotoxin Quant Kit (Thermo Fischer Scientific, Rockford, IL). A high endotoxin calibration curve (0.1-1.0 EU/mL) was used to determine the endotoxin concentration.

### HIV-1 Gp120 ELISA binding assays

Purified, recombinant HIV-1 Gp120 was immobilized on high-binding ELISA plates (Greiner #655081). Plates were washed with PBS-T and blocked with a solution of 5% (w/v) bovine serum albumin (BSA, Fisher #BP9706-100) in 1X PBS pH 7.4 (PBS). For evaluating the binding of GRFT samples to gp120, plates were washed three time with PBS-T and incubated with serial half-log dilutions of GRFT, diluted in PBS, for one hour at room temperature. Plates were then washed three times with PBS-T and incubated with rabbit anti-GRFT polyclonal antibodies for one hour at room temperature. Plates were again washed three times with PBS-T and incubated with goat anti-rabbit IgG-HRP conjugate (Thermo Fisher Scientific #31460) for one hour at room temperature. Plates were finally washed three more times with PBS-T and developed using 1-Step Ultra TMB-ELISA solution (Thermo Fisher Scientific #34028). The HRP reaction was stopped with 1M hydrochloric acid and absorbance values at 450 nm were measured on a SpectraMax i3x plate reader (Molecular Devices).

### Anti-SARS-CoV-2 pseudovirus assay

HEK-293T/17 cells (ATCC) were maintained in Dulbecco’s modified Eagle’s medium (DMEM) with 10% fetal bovine serum, 100 IU/mL penicillin and 100 μg/mL streptomycin (Thermo Fisher Scientific). HEK-293T cells overexpressing human ACE2^47^ were obtained from BEI Resources and maintained in DMEM as above. SARS-CoV-2 pseudovirus particles encoding for luciferase expression were prepared using the plasmids and protocols essentially as previously described^48^. The pseudoviral infectivity assay was also similar to previous publications^48^. Briefly, HEK293-ACE2 cells were seeded in 384-well white plates (PerkinElmer, USA), allowed to attach overnight followed by addition of pseudoviral particles in the presence or absence of varying concentrations of GRFT. After 3 d incubation, luciferase reagent was added (PerkinElmer, USA) and expression was quantitated (Spectrostar Omega plate reader – BMG Labtech). After normalization (absence of GRFT = 100%), dose-response curves were generated and IC50 values estimated by 4-parameter logistic analysis using GraphPad Prism software.

## Results

### Expression of Recombinant GRFT using Cell-free systems

Initial *E. coli* CFPS was first evaluated with a gene (tagless-GRFT.std) consisting of the same frequently used codon for each amino acid. Disappointingly low yields (≈ 200μg/ml) and product solubility (≈50%) resulted despite evaluating various expression temperatures, incubation times, amino acid concentrations, ionic strengths, and chaperone additions. Recognizing that GRFT has an abnormally high frequency of glycine’s (18.2%) and serine’s (15.7%), a new gene was designed to randomize the use of five of the six serine codons (AGC was not used) and all four of the glycine codons. Since these codons are GC-rich, the gene was further modified to minimize the stability of potential mRNA secondary structure. The optimized GRFT gene (tagless-GRFT.opt) provided a 3.8-fold improvement in total protein accumulation and a 6.3-fold improvement in soluble product (Figure 2). Autoradiography indicated accumulation of the full-length 12.7 kDa GRFT without any visible truncated products (Supplementary Figure 2). Due to the high abundance of glycine and serine codons within the gene, the variability in codon usage for these amino acids within the optimized gene probably enabled a more consistent supply of amino acylated glycine and serine tRNAs. It is also possible that secondary structure avoidance was beneficial. Addition of 2 mM additional glycine and serine was not helpful. Additionally, reducing the potassium glutamate concentration from 175 to 50 mM did not improve protein folding, and utilization of a BL21-sourced extract was not helpful (Figure 2). The total protein yield was about 900 μg/mL and about 80% was soluble (Figure 2A).

**Figure 2.**
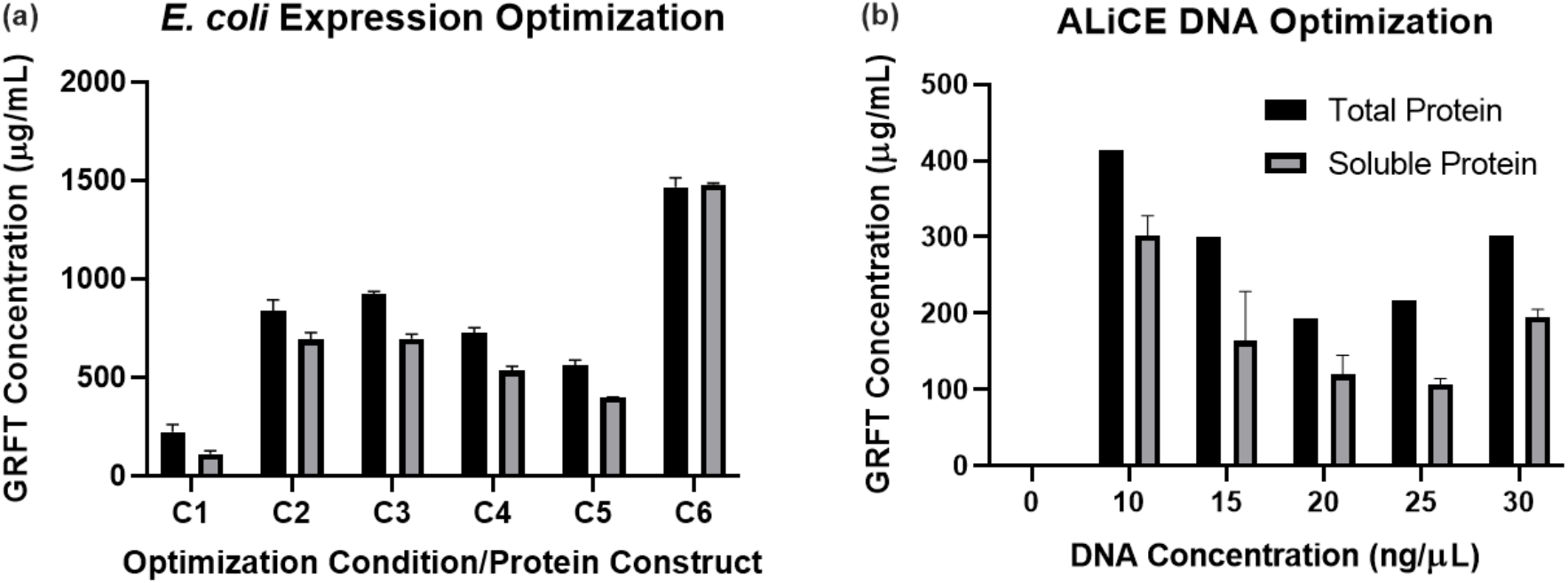
**(a)** GRFT yields from an *E. coli* CFPS system comparing two different codon-optimized constructs, tagless-GRFT.std and tagless-GRFT.opt. Reactions were performed in triplicate at a 25 μL scale overnight at 30 °C. C1, tagless-GRFT.std; C2, tagless-GRFT.opt; C3, tagless-GRFT.opt + 2 mM Serine/Glycine; C4, tagless-GRFT.opt with 50 mM KGlu (normal is 175mM); C5, tagless-GRFT.opt using BL21 Extract; C6, sfGFP control. **(b)** Tagged-GRFT yields from various plasmid concentrations in pALiCE01 expression vector. Reactions were performed in triplicate at the 250 μL scale overnight at 25 °C. Total GRFT concentration was determined in pooled samples. Error bars represent standard deviation (SD) of the mean (n=3).

In comparison, the ALiCE^®^ system using plant (*N. tabacum* BY-2*J* cell extracts was evaluated using both the pALiCE01 plasmid for cytosolic accumulation and pALiCE02 plasmid for microsomal accumulation. Even though GRFT has no disulfide bonds, the latter was evaluated to assess any potential benefit from microsomal translocation or chaperones. No product accumulation was detected using the pALiCE02 plasmid, and the pALiCE01 plasmid produced a maximum total protein yield of 410 μg/mL for the 14.50 kDa tagged-GRFT product. Surprisingly, best yields were obtained with a DNA concentration of 10 ng/uL and higher concentrations were harmful (Figure 2B). Approximately 75% of the total product was soluble after centrifugation of the ALiCE^®^ harvested fluid.

### IMAC and Heat-induced precipitation purification

The tagged-GRFT construct was purified using the IMAC methods described earlier resulting in a recovery of 36% after a 0.5 column volume, 250mM imidazole elution (Figure 3a). Approximately, 33% of the tagged-GRFT mixture was collected in the flow-through (FT) fraction suggesting incomplete capture, but less than one percent was collected in the Wash 1(W1) step (Figure 3a). Alternatively, by using previously optimized heat-induced purification conditions by Decker et al.^28^ (20.5% saturation (NH4)2SO4, pH 3.4, 60°C for 1h) we obtained 50% and 55% recovery of soluble tagged- and tagless-GRFT, respectively (Figure 3b).

**Figure 3.**
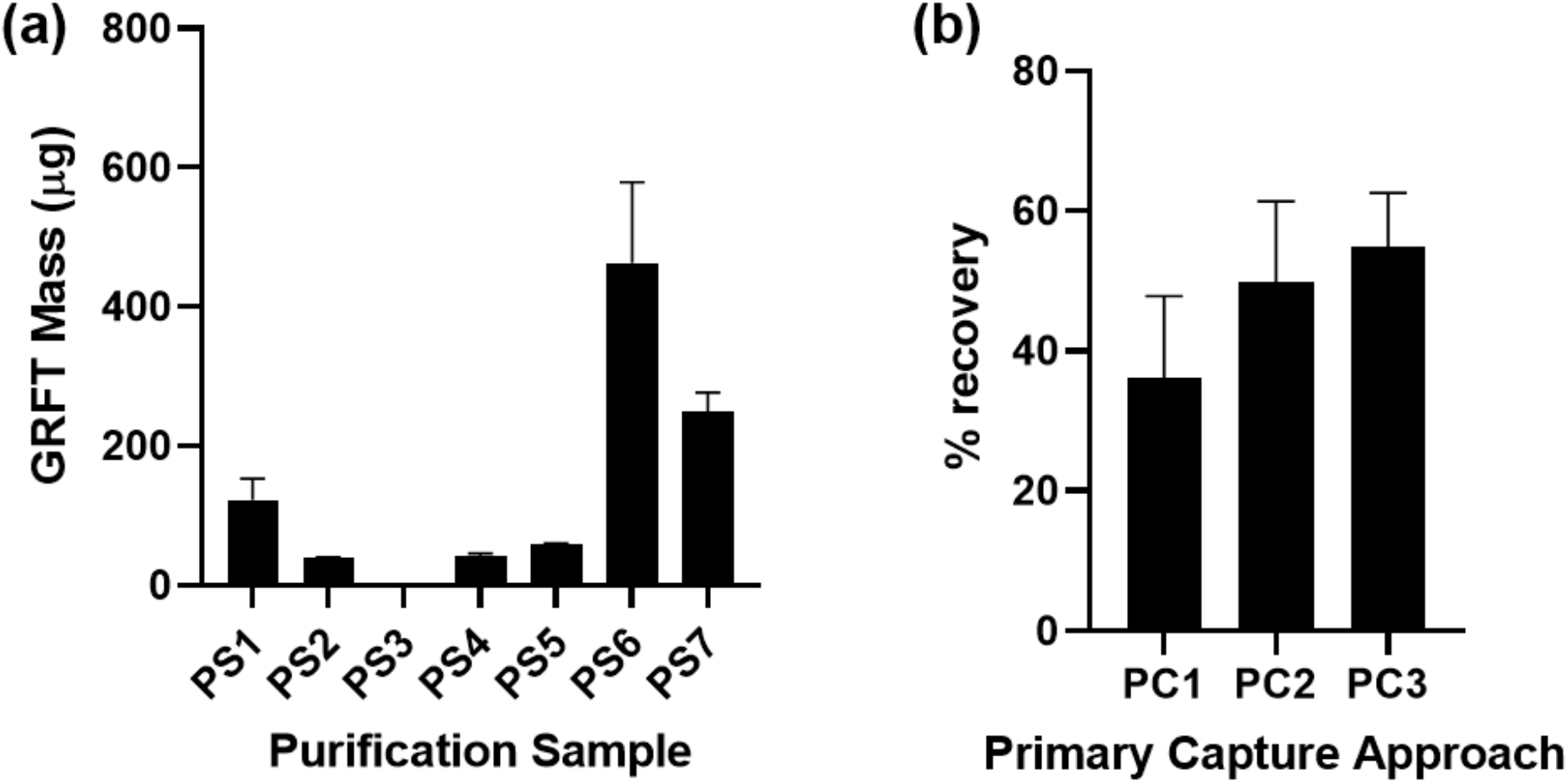
(a) GRFT amounts in various harvest and purification fractions as estimated by Western blotting analysis (Supplementary Figure 3) in the primary capture step of GRFT purification for tagged- and tagless-GRFT constructs expressed in ALiCE^®^ and *E. coli* cell-free systems, respectively. PS1, tagged-GRFT Harvest; PS2, tagged-GRFT IMAC FT; PS3, tagged-GRFT IMAC W1; PS4, IMAC Elution; PS5, tagged-GRFT Post Precipitation; PS6, tagless-GRFT.opt Harvest; PS7, tagless-GRFT.opt Post Precipitation. PS1-7 were then utilized in a mass balance to determine protein recovery. (b) (%) recovery obtained for IMAC (tagged-GRFT, PC1) and heat-induced purification (tagged-GRFT, PC2 and tagless-GRFT.opt, PC3). Error bars represent SD of the mean (n=2).

### Size exclusion chromatography (SEC) on recombinant GRFT

GRFT typically is recovered as a homodimer with a somewhat variable hydrodynamic radius dependent upon the degree of domain swapping in the tertiary structure. Essentially complete homodimerization was confirmed using SEC. Molecular weight estimates for tagged GRFT were 31.2 and 29.8 kDa for samples purified using IMAC (Supplementary Figure 4a) and heat precipitation (Supplementary Figure 4b), respectively in reasonable agreement with but larger than the expected value of 29,060 kDa. The SEC estimate for tagless-GRFT purified using heat precipitation was 24.2 kDa, smaller than the actual molecular weight of the dimer, 25,376 kDa (Supplementary Figure 4c). While the SEC estimates are within expected experimental error, they suggest that the purification tag may have caused a larger than expected tertiary structure, possibly by affecting domain swapping in the dimer.

### Molecular weight determination of recombinant GRFT using HPLC-TOF-MS

HPLC-TOF analysis of the products expressed in both *E. coli* and ALiCE^®^ cell-free systems indicated molecular weights in conformity with those expected. The HPLC-TOF mass spectrum of tagless-GRFT.opt, tagged-GRFT, and NIH GRFT standards yielded +1 mass ion species of 12,688 Da, 14,530 Da, and 14,499, respectively (Figure 4a-c). The theoretical molecular weight (Benchling, USA) assuming N-terminal methionine removal are obtained to be 12,687 Da, 14,504 Da, and 14,499 Da. for each sample, respectively. The reason for the tagged-GRFT discrepancy is not known.

**Figure 4.**
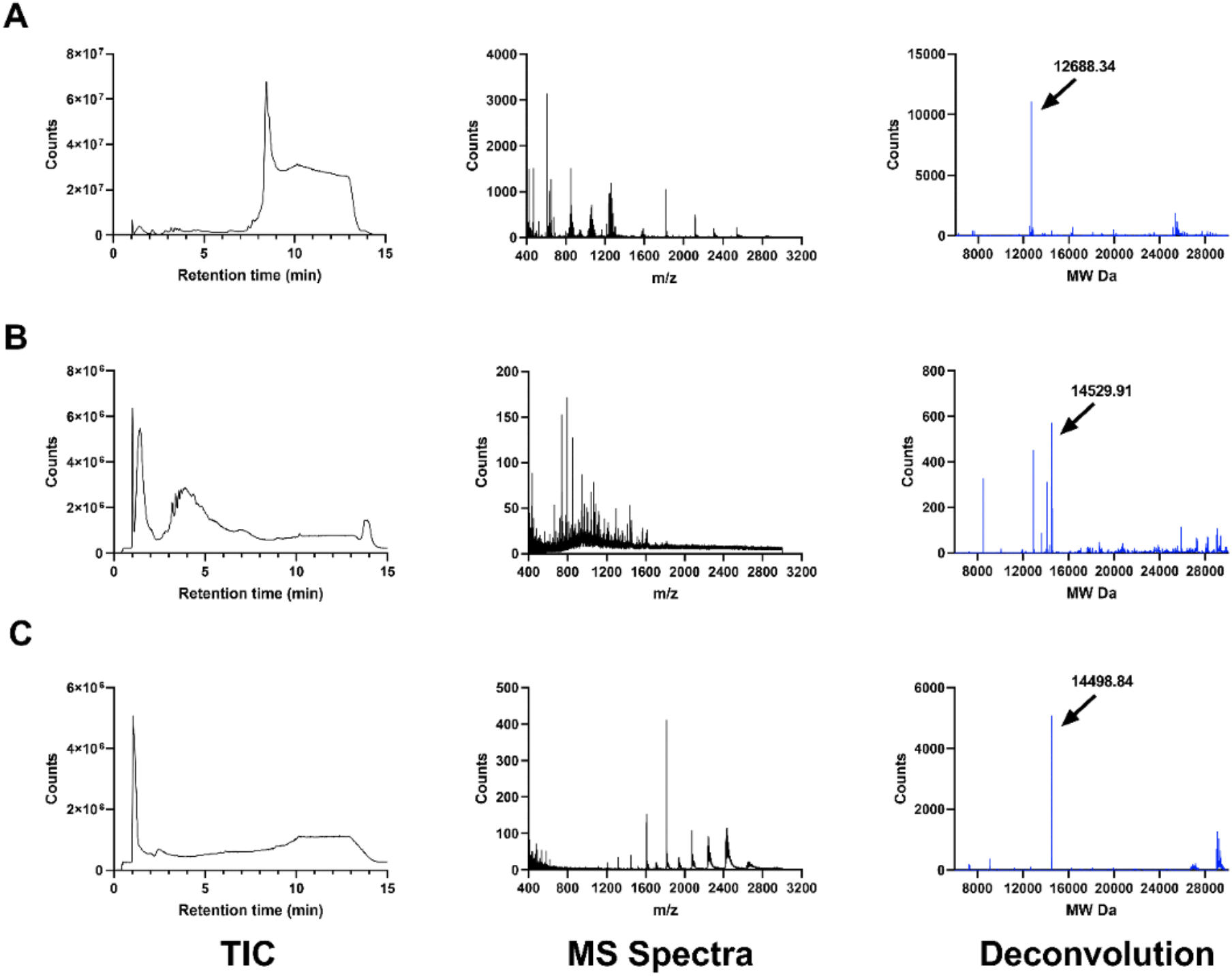
**LC/MS spectra** Total Ion Chromatogram (TIC), MS spectrum, and deconvolution of *E. coli* and ALiCE^®^ post precipitation products in sterile-filtered 1X PBS (pH 7.4). (A) *E. coli* expressed tagless-GRFT.opt (MW 12.68 kDa) (Left-top). (B) ALiCE^®^ expressed tagged-GRFT (MW 14.53 kDa) (Middle) (C) NIH GRFT standard (MW 14.49 kDa) (Right-bottom).

### AEX Chromatography and Endotoxin Quantification

SDS PAGE analysis of samples collected after loading the tagless-GRFT into the AEX column revealed no significant attraction to the HiTrap™ Q FF resin (Supplementary Figure 5). The resulting AEX FT fraction had about a 13-fold reduction in endotoxin concentration. The indicated concentration of about 0.2 EU/ml (Supplementary Figure 6) is well below the 0.5EU/mL limit indicated by the FDA for nasally administered agents. Furthermore, the purity of RT fractions collected from 0.25 to 1.50 min was assessed as >99% by SDS-PAGE/ImageJ analysis (Supplementary Figure 7) with a final estimated concentration of (0.065 ± 0.012 mg/mL).

### Functional Assessment of *E. coli* and ALiCE^®^ produced GRFT Samples

To determine if the *E. coli* and ALiCE^®^ CFS produced GRFT samples were functionally active against viruses, we evaluated both envelope glycoprotein binding (HIV-1 gp120) and *in vitro* inhibition of viral entry (SARS-CoV-2). First, samples were tested for their ability to bind to the HIV-1 envelope glycoprotein gp120. This assay has been used previously for quality control of GRFT samples used in clinical trials^21^. As can be seen in Figure 5A, CFS production of GRFT resulted in protein with binding affinity (≈30nM) to gp120 that was similar to GRFT produced previously in *E. coli* ^49^. This result indicates that both *E. coli* and ALiCE^®^ CFS can produce biologically active GRFT. To further characterize the biological activity of the GRFT produced in CFS, GRFT samples were evaluated for their inhibition of SARS-CoV-2 pseudoviral binding to ACE2 positive mammalian cells. As shown in Figure 5B, both samples of GRFT produced in cell-free systems inhibited the infection of mammalian cells by SARS-CoV-2. The IC50 value for tagged-GRFT was calculated to be 510 nM and that of tagless-GRFT.opt was determined to be 324 nM, similar to those previously reported^27^. Further testing of both cell-free GRFT samples for mammalian cell cytotoxicity indicated that neither sample caused any loss of cell viability at the highest tested concentrations.

**Figure 5.**
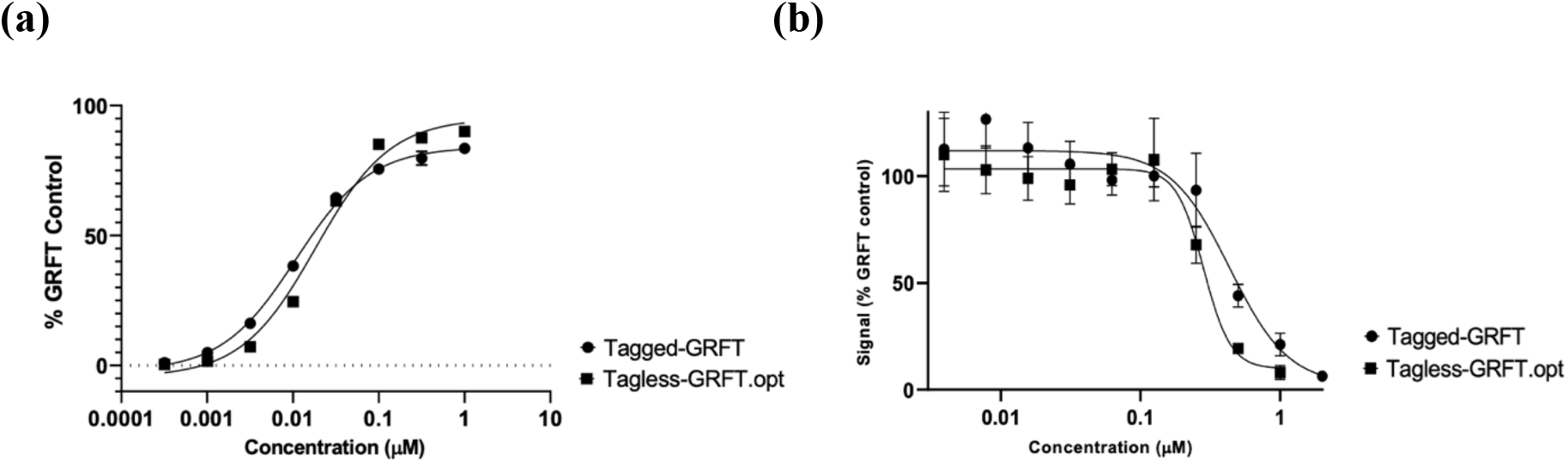
(a) Side by side comparison of gp120 ELISA for Tagged-GRFT (ALiCE^®^) and tagless-GRFT.opt (*E. coli*) CF expressed products while evaluating binding at varied concentrations of GRFT. (b) Activity of GRFT in SARS-CoV-2 pseudovirus assays showing inhibition of pseudoviral entry. Samples tested in duplicate, and experiment repeated twice. Visualization using luciferase endpoint. Error bars represent SD of the mean (n = 4).

## Discussion

The ability to produce broad spectrum antiviral biologics in a rapid, scalable fashion at the site of pandemic emergence could provide critical advantages for suppressing future pandemics. To address this opportunity, we report cell-free bioprocess options for the production and purification of a broad-spectrum antiviral, GRFT, with the potential for hindering viral transmission. Our selection of CFSs was motivated by its scalability, portability, and ability to stockpile all major reagents in a dried state, ready for instant activation. Potential POC settings include pharmacies, rural hospitals, physician’s offices, and field medical units designed for underserved regions. Additionally, we expect that this opportunity will motivate academic labs, particularly those in structural and synthetic biology, to help generate novel anti-viral proteins that are even more effective.

Here, we show that both the *E.coli* and ALiCE^®^ (*N. tabacum*) based CFPS systems are capable of synthesizing active GRFT in under 24 hours. We first chose to explore an *E. coli*-based CFPS system due to its extensive optimization over the last 20 years. Feasibility was suggested by *in vivo E.coli* GRFT production^19,49^. However, initial yields were surprisingly low. We realized that glycine and serine residues were highly abundant in the protein and that the initially designed gene had used the same GC rich codon for each amino acid. Randomizing codon usage and decreasing the stability of predicted mRNA secondary structure provided impressive improvements. The newly designed variability in codon usage for these amino acids probably enabled a more consistent supply of amino acylated glycine and serine tRNAs. It is also possible that secondary structure avoidance was beneficial. Future work will seek to evaluate the relative contributions of each influence. The tagless-GRFT.opt gene increased the soluble yield to 700-900 μg/ml. Since the expression of GFP (Figure 2) was nearly 2-fold higher, it is possible that additional gene optimization could be beneficial.

Alternatively, given reports of successful expression of GRFT in tobacco-based cell lines, we evaluated the ALiCE^®^ cell-free expression platform as an analogous cell-free approach to the current *in vivo* approaches^45,50^. Initially, we opted to utilize the pALiCE02 vector to enhance protein folding but observed little to no protein expression. For this reason, the tagged-GRFT construct was subcloned into the pALiCE01 vector for cytosolic expression, resulting in substantially improved yields. Assessing the effects of increasing plasmid DNA concentration from 10 to 30 ng/μL revealed that the lowest final DNA concentration, 10 ng/μL, yielded the highest soluble protein yield (302 μg/mL). Remarkably, an increase in DNA concentration did not enhance protein titer. This is likely due to the fixed (unknown) concentration of magnesium ions present in the ALiCE^®^ reaction mixture which will partially complex with negatively charged components in the cell free reaction, such as DNA^51^, resulting in non-optimal Mg^++^ availability. Additionally, we believe that further codon optimization utilizing a similar method to those utilized for *E. coli* mentioned above, may also be beneficial in improving yield in the ALiCE system.

As previously reported, we sought to leverage the unique thermostability and acid stability of GRFT in an effort to display whether similar behavior could be observed when purifying material from a cell-free reaction product. Here, we report a GRFT recovery of approximately 55% when utilizing the heat-induced precipitation purification rather than conventional IMAC methods (36% yield). Heat-induced purification and AEX chromatography, yielded tagless-GRFT with a purity of greater than 99% with an endotoxin concentration below the FDA limit of 0.5 EU/mL. This product was then analyzed further for its identity and activity, confirming anti-HIV and anti-SARS-CoV-2 activity nearly indistinguishable from that of GRFT expressed *in vivo*.

Furthermore, the rapid expression and purification of GRFT mentioned here can be readily implemented on the Bio-MOD platform developed by Adiga et al.^36^. The current automated Bio-MOD platform has two modules: the protein expression module and the protein purification module, each with associated analytics. The system is fully automated with built-in software and programmable syringe pumps with pressure sensors for the delivery of lysate and buffers. Protein expression is currently carried out in dialysis cassettes. Once requisite components (cell lysate, reaction mix and cDNA for the target protein) are reconstituted, the cassette is then immersed in dialysis buffer inside a sealed bag. The current onboard protein expression module is equipped with a Peltier heating source, temperature sensor, and muffin-fan for a uniform and continuous distribution of heat. Both the temperature and shaking speed are programmable via LabVIEW software program using a computer tablet. The protein purification module has two built-in flow cells, each equipped with a UV sensor for monitoring the 2-step purification process involving affinity chromatography and ion-exchange chromatography for polishing. In addition, four pressure sensors are incorporated behind each syringe pump plunger for continuous system pressure monitoring. These along with system temperature sensors form the current process analytical technologies (PAT) that are already proving their value in demonstrating process consistency^52^. For the present application a simple modification that raises the expression reactor temperature to 60°C in addition to a syringe pump to infuse ammonium sulfate solution into the reactor is all that is required and can be readily implemented in the place of the current affinity purification step. After heat treatment, supernatant can be removed after the precipitated proteins have settled to the bottom of the bioreactor. If needed, simple filtration can further clarify the purified product containing solution.

By using heat-induced purification, downstream processing is made easier and more cost efficient given the absence of costly resins, buffers, and equipment typically used during IMAC purification. Techno-economic analysis by Decker et al. have shown that using the heat-induced purification approach leads to a 6-fold reduction in downstream purification capital expenses in comparison to conventional chromatography methods^28^. While a technoeconomic model has yet to be performed on the cell-free expression of GRFT, optimization of cell-free reagents has shown a 95 percent reduction in CFPS cost, bringing the cost per liter of a cell-free mastermix to approximately 80 USD^53^. Additionally, the cost per liter of *E.coli* lysate can be approximated to be 20 USD based on previous techno-economic analysis^54^. Furthermore, assuming downstream processing to be 40 percent of the total bioprocessing cost, this would mean a 3 mg daily dose of GRFT^34^ (assuming 0.5g/L protein recovery) would cost roughly 0.85 USD, making it accessible to a global patient population. In comparison, previous in-silico analysis by Alam et al. estimate a cost of approximately 0.32 USD per dose of GRFT^44^. This is significantly less than currently available antivirals, such as Paxlovid, which cost upward of 100USD per day. Additionally, while other antiviral proteins, such as mAbs are available for treatment of COVID-19, these proteins require expensive Protein A/G resins and buffers during purification, likely contributing to their exorbitant price^46,55^. More notably, while mAbs have the highest safety and efficacy profile among biologics^56,57^, their reduced efficacy against SARS-CoV-2 variants is an indication that a stronger and more broad-spectrum antiviral such as GRFT could serves as a better alternative in suppressing the spread of rapidly emerging SARS-CoV-2 variants^58^.

As previously shown, lyophilized cell extracts have been used to produce many therapeutic proteins^36,52^. Having the operational capacity for rapid distributed manufacturing could be vital infrastructure in dealing with future viral threats as well as other health challenges that require biologics, including for example, a radiation exposure event^52^. Through integrating the bioprocess in this study with an automated biomanufacturing platform such as that developed by Adiga et al.^36^, the decentralized manufacturing of GRFT would be made more achievable and will allow for a targeted and effective response in areas of the world where distribution of antivirals is challenging. We also note that the technology approach proposed here is generally applicable. Many biologics can be manufactured using the same approach. Given the worldwide threat of future viral outbreaks, investments in such mitigation approaches are required now.

## Author Contribution

**Experiment Design and Data Analysis**

S.B., M.L., J.S., B.O., G.R.

**Griffithsin Gene Construction and DNA preparation**

M.L., S.B.

**E. coli Cell Extract Preparation**

M.L.

**E. coli Cell-free Expression and Quantification**

M.L.

**Autoradiogram Preparation**

M.L

**ALiCE^®^ Expression**

S.B.

**Heat Precipitation of tagless and tagged GRFT**

S.B.

**Immobilized Metal Affinity Chromatography (IMAC) capture of tagged-GRFT**

S.B.

**Western Blot Analysis**

S.B.

**Coomassie and Silver Staining**

S.B

**SEC Analysis**

A.T., S.B.

**HPLC-TOF analysis**

D.T.

**Anion Exchange Chromatography and Endotoxin Removal**

A.T. S.B.

**Gp120 Assay/Kinetic**

B.O., L.K., J.W., C.H.

**Anti-SARS-CoV-2 assay**

B.O., L.K., J.W., C.H.

**Material Distribution and Project Management**

S.B. G.S.S., B.O., D.L., A.G., V.M., C.L., D.F., J.S. G.R.

**Editing**

All Individuals

## Acknowledgements

Mike Tolosa for helping generated a schematic of our distributed manufacturing platform. LenioBio for providing ALiCE Lysate.

NIH/NCATS for material support.

This project has been funded in whole or in part with Federal funds from the Frederick National Laboratory for Cancer Research, National Institutes of Health, under contract HHSN261200800001E. The content of this publication does not necessarily reflect the views or policies of the Department of Health and Human Services, nor does mention of trade names, commercial products or organizations imply endorsement by the US Government

