## Supplemental Figure 1-7 for "An approach to rapid distributed manufacturing of broad spectrum anti-viral griffithsin using cell-free systems to mitigate pandemics"

### Supplementary Documents

**tagged-GRFT**
ATGGGGAGCTCTCATCACCACCACCATCATTCTTCTGATGACGATGATAAATCACTGACTCATAGAAGTTTGGAGGA
AGTGGGGGGTCTCCCTTTTCAGGCCTTTCCTCCATTGCTGTACGAAGTGGCTCATACTTGGATGCCATCATAATAGAT
GGGGTTCATCATGGAGGCAGTGGTGGAAACTTGTCTCCAACTTTCACTTTTGGTTCTGGTGAATATATTTCAAATATG
ACAATTCGTTCGGGAGATTACATTGATAATATATCATTTGAGACAAATCAGGGAAGGAGATTCGGTCCTTATGGTGGT
TCAGGAGGCAGCGCAAACACCCTCAGCAATGTGAAAGTTATTCAAATCAATGGATCTGCTGGTGATTATCTTGACAG
TTTAGACATTTACTATGAACAATATTGA

**tagless-GRFT.opt**
ATGTCATTAACTCATAGAAAATTTGGTGGTTCCGGTGGCTCTCCGTTTAGTGGGCTGTCCAGTATTGCGGTGCGCTCC
GGGTCATATCTGGATGCGATTATTATTGATGGGGTGCATCATGGTGGCTCTGGTGGTAACCTGTCTCCGACCTTTACCT
TTGGTTCAGGTGAATATATTTCTAACATGACCATTCGCTCGGGTGATTATATTGATAACATTTCATTTGAAACCAACCA
GGGTCGCCGCTTTGGTCCGTATGGTGGCTCTGGTGGCTCTGCGAACACCCTGTCGAACGTGAAAGTGATTCAGATTA
ACGGTTCTGCGGGCGATTATCTGGATTCACTGGATATTTATTACGAACAGTATTAA

**Supplementary Figure 1a. tagged-GRFT and tagless-GRFT.opt DNA Sequence**
DNA sequence of recombinant tagged-GRFT and tagless-Q-GRFT construct.

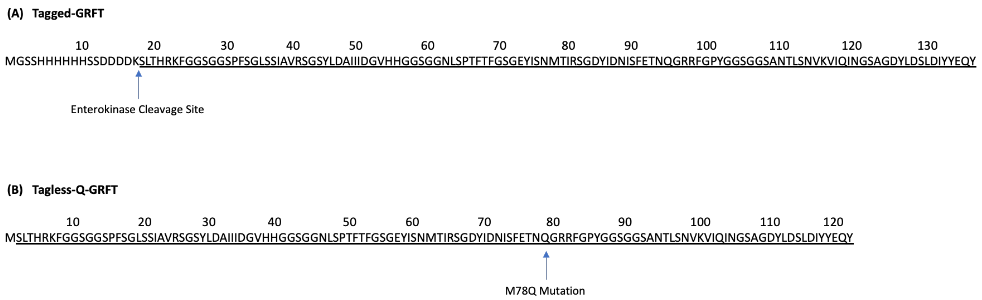


**Supplementary Figure 1b. tagged-GRFT and tagless-GRFT Amino Acid Sequence**
Amino acid sequence of recombinant tagged-GRFT and tagless-Q-GRFT construct. (A) tagged-GRFT insert (wild-type) with a N-terminal extension containing a hexa-histidine tag and enterokinase cleavage site for tag removal. (B) tagless-Q-GRFT insert which contains amino acid substitution at position 78 from methionine (M) to glutamine (Q), in order to reduce oxidation of expressed protein. Underlined sequences indicate the target GRFT expressed in ALiCE and *E.coli*, respectively.

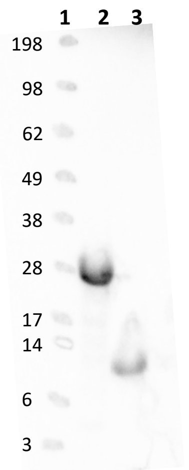


**Supplemental Figure 2. Autoradiogram of tagless-GRFT.opt**
Soluble portions of overnight, 37ºC, *E. coli* CFPS of sfGFP (Lane 2) and tagless-GRFT.opt (Lane 3). Experiment was performed in quadruplicate, pooled, and 10ul was loaded onto a 4-12% SDS gel that was then dried and transferred to a phosphor screen for autoradiography. Autoradiography indicates incorporation of 14C-leucine into the synthesized peptide. SeeBlue™ Plus 2 pre-stained ladder was loaded into Lane 1 at 6ul and spotted with 14C-leucine.

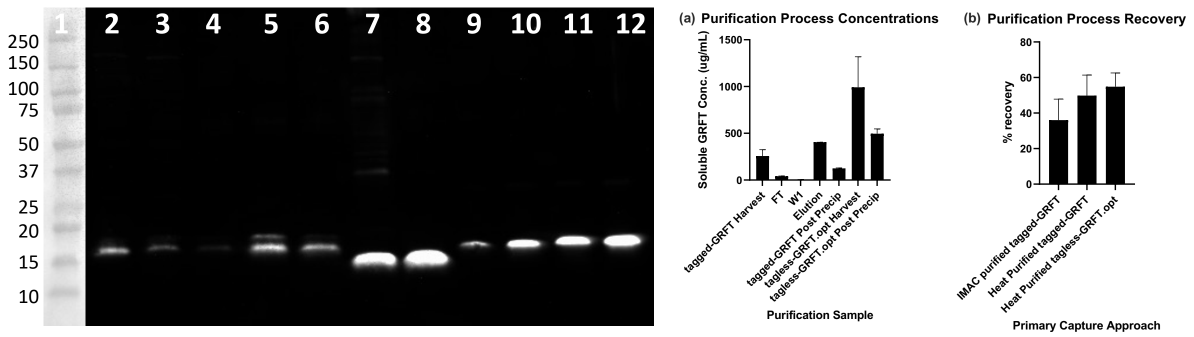


**Supplementary Figure 3. Western blot analysis (probed w/ anti-GRFT antibody) for determination of protein recovery during purification** Lane 1, Marker; Lane 3-5 samples from tagged-GRFT IMAC purification; Lane 2, tagged-GRFT Harvest (1.5µL); Lane 3, FT (3µL); Lane 4, W1 (10µL); Lane 5, 100µL Elution (3µL); Lane 6, tagged-GRFT Post Precipitation (3µL); Lane 7, tagless-GRFT.opt Harvest (1.5µL); Lane 8, tagless-GRFT Post Precipitation (3µL); Lane 9-12, 250, 500, 750, 1000ng GRFT Standard, respectively. Error bars represent standard deviation of the mean (n=2).

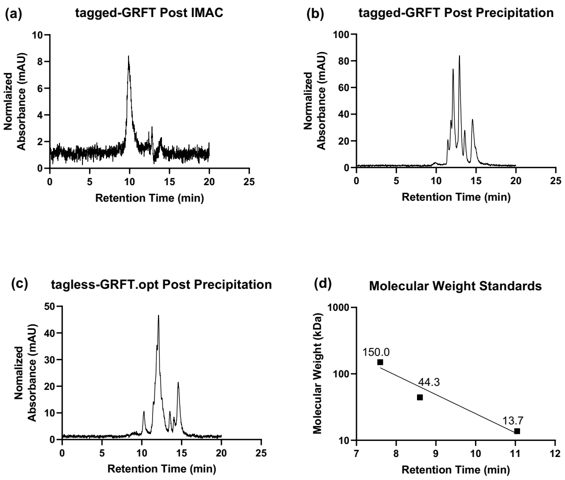


**Supplementary Figure 4.** Chromatograms showing GRFT after primary capture purification(a) SEC run using 50 µL injection of IMAC purified tagged-GRFT. (b) SEC run using 10 µL injection of the tagged-GRFT post precipitation. (c) SEC run using 10 µL injection of the tagless-GRFT.opt post precipitation. (d) Standard linear regression generated from protein standard mix whose molecular weights are within the inclusion range of the SEC column allowing for calculation of molecular weight of GRFT samples.

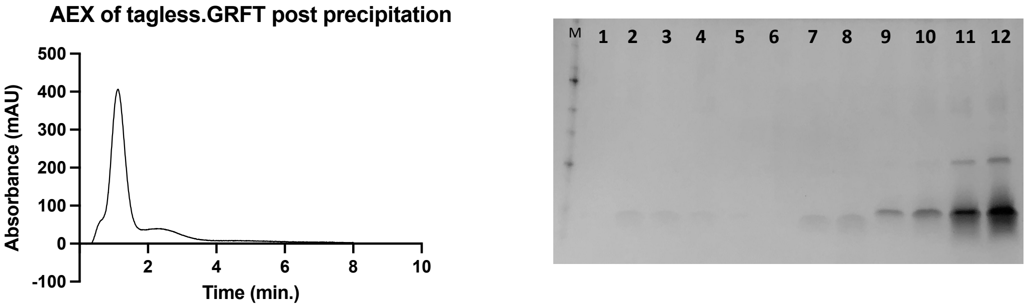


**Supplementary Figure 5.** AEX chromatograms (Left) from a 125µL injection of tagless-GRFT.opt post precipitation sample into a 1 mL HiTrap™ Q FF column in flow through mode (1mL/min). Samples were manually collected every 0.25 min till return to baseline (~1.5min). Silver stain SDS-PAGE (Right) of RT fractions 0-0.25, 0.25-0.50, 0.50-0.75, 0.75-1.0, 1.25-1.50, 1.50-1.75, lane 1-6, respectively; Pooled AEX samples, RT fractions 0.25-1.50 (10 and 20uL), lanes 7-8, respectively; 250, 500, 750, 1000ng GRFT standard, lanes 9-12, respectively.

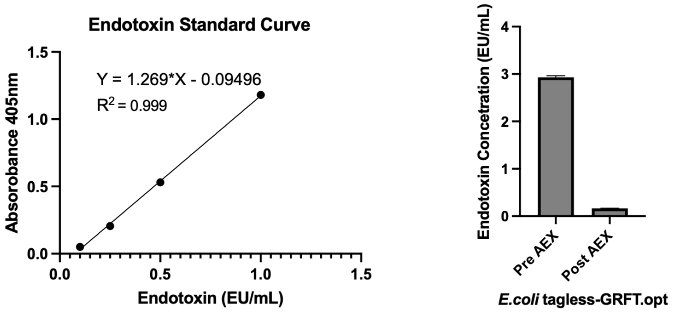


**Supplementary Figure 6. LAL chromogenic assay for endotoxin quantification** AEX sample from peak fraction (RT, 0.25 - 0.50) run on Pierce Endotoxin Quant Kit. A high standard curve (0.1 -1EU/mL) was used to assess the concentration of pre and post AEX endotoxin concentrations for the tagless-GRFT.opt construct.

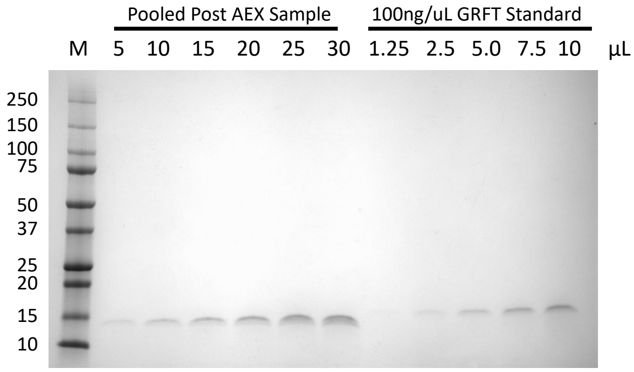


**Supplementary Figure 7. Coomassie R-250 Staining** Purity was calculated to be > 99% by ImageJ densitometry, with a final concentration of 0.0651 ± 0.0116 mg/mL
